# DCI: Learning Causal Differences between Gene Regulatory Networks

**DOI:** 10.1101/2020.05.13.093765

**Authors:** Anastasiya Belyaeva, Chandler Squires, Caroline Uhler

## Abstract

**Summary:** Designing interventions to control gene regulation necessitates modeling a gene regulatory network by a causal graph. Currently, large-scale expression datasets from different conditions, cell types, disease states and developmental time points are being collected. However, application of classical causal inference algorithms to infer gene regulatory networks based on such data is still challenging, requiring high sample sizes and computational resources. Here, we propose an algorithm that efficiently learns the differences in gene regulatory mechanisms between different conditions. Our difference causal inference (DCI) algorithm infers changes (i.e., edges that appeared, disappeared or changed weight) between two causal graphs given gene expression data from the two conditions. This algorithm is efficient in its use of samples and computation since it infers the differences between causal graphs directly without estimating each possibly large causal graph separately. We provide a user-friendly Python implementation of DCI and also enable the user to learn the most robust difference causal graph across different tuning parameters via stability selection. Finally, we show how to apply DCI to bulk and single-cell RNA-seq data from different conditions and cell states, and we also validate our algorithm by predicting the effects of interventions.

**Availability and implementation:** All algorithms are freely available as a Python package at http://uhlerlab.github.io/causaldag/dci

## 1 Introduction

Biological processes from differentiation to disease progression are governed by gene regulatory networks. Over the past few decades, various methods have been developed for inferring gene regulatory networks from gene expression data (Wang and Huang, 2014). The majority of methods learn undirected graphs of interactions between genes such as correlation-based coexpression networks (Langfelder and Horvath, 2008), Gaussian graphical models that capture partial correlations (Friedman *et al.*, 2008), or networks that measure dependencies between genes using mutual information (Reshef *et al.*, 2011). However, the ultimate goal is often to use gene regulatory networks to predict the effect of an intervention (small molecule, overexpression of a transcription factor, knock-out of a gene, etc.). This cannot be done using an undirected graph and necessitates modeling a gene regulatory network by a causal (directed) graph.

One of the most common frameworks for representing causal relationships are directed acyclic graphs (DAGs). A variety of methods including the prominent PC and GES algorithms have been developed for learning causal graphs from observational data (Glymour *et al.*, 2019). These methods have been successfully applied to learning (directed) gene regulatory networks on a small number of genes, starting with a pioneering study by Friedman *et al.*, 2000. However, applying these methods at the whole genome-level is still challenging due to high sample size and computational requirements of the algorithms.

We address this problem by noting that it is often of interest to learn *changes* in causal (regulatory) relationships between two related gene regulatory networks corresponding to different conditions, disease states, cell types or developmental time points, as opposed to learning the full gene regulatory network for each condition. This can reduce the high sample and computational requirements of current causal inference algorithms, since while the full regulatory network is often large and dense, the difference between two related regulatory networks is often small and sparse. As of now, this problem has only been addressed in the undirected setting, namely by KLIEP (Liu *et al.*, 2017), DPM (Zhao *et al.*, 2014) and others (Fukushima, 2013; Lichtblau *et al.*, 2017) that estimate differences between undirected graphs; for a recent review see Shojaie (2020).

With the recent advances in high-throughput single-cell RNA-sequencing under different contexts and cell types (Zheng *et al.*, 2017), there is a growing need for methods that learn changes between gene regulatory networks. Causal inference algorithms that infer changes between gene regulatory networks could for example reveal that a particular gene controls different sets of target genes in different conditions. While for small number of genes, it is feasible to apply current causal structure discovery algorithms to learn the causal graphs for each condition seperately and take the difference of the graphs, this approach is highly inefficient in its use of samples and computation. In this paper, we describe the *difference causal inference* (*DCI*) algorithm for direct estimation of the difference causal graph based on observational data from two conditions. For theoretical properties of this algorithm, in particular the proof that DCI provides consistent estimates of the difference causal graph, and simulations showing that it outperforms the naive approach of separate estimation of each causal graph and subsequent computation of the difference, see Wang *et al.* (2018). In this paper, we present an easy to use Python package to apply DCI to gene expression data from different conditions and demonstrate the algorithm’s performance on predicting the effects of interventions on single-cell RNA-seq data. Importantly, our DCI Python implementation also allows selecting the most robust difference gene regulatory network based on a collection of tuning parameters via stability selection (Meinshausen and Bühlmann, 2010). We also include a tutorial and an example use case of DCI on a bulk RNA-seq dataset from two ovarian cancer patient cohorts with different survival rates from Tothill *et al.* (2008) at http://uhlerlab.github.io/causaldag/dcLtutorial To seamlessly integrate DCI with other causal inference methods, we incorporated our DCI code into the larger `causaldag` package.

## 2 Difference Causal Inference (DCI) package

DCI takes as input two gene expression matrices 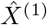 and 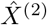 of size *n*_1_ × *p* and *n*_2_ × *p*, where *n*_1_ and *n*_2_ denote the number of samples in each dataset and *p* denotes the number of genes. These matrices contain the RNA-seq values corresponding to two different conditions, such as healthy and diseased, different cell types, or different time points. DCI outputs the difference causal graph between the two conditions, i.e. the edges in the gene regulatory networks that appeared, disappeared or changed weight between the two conditions (Fig. 1).

**Figure 1:**
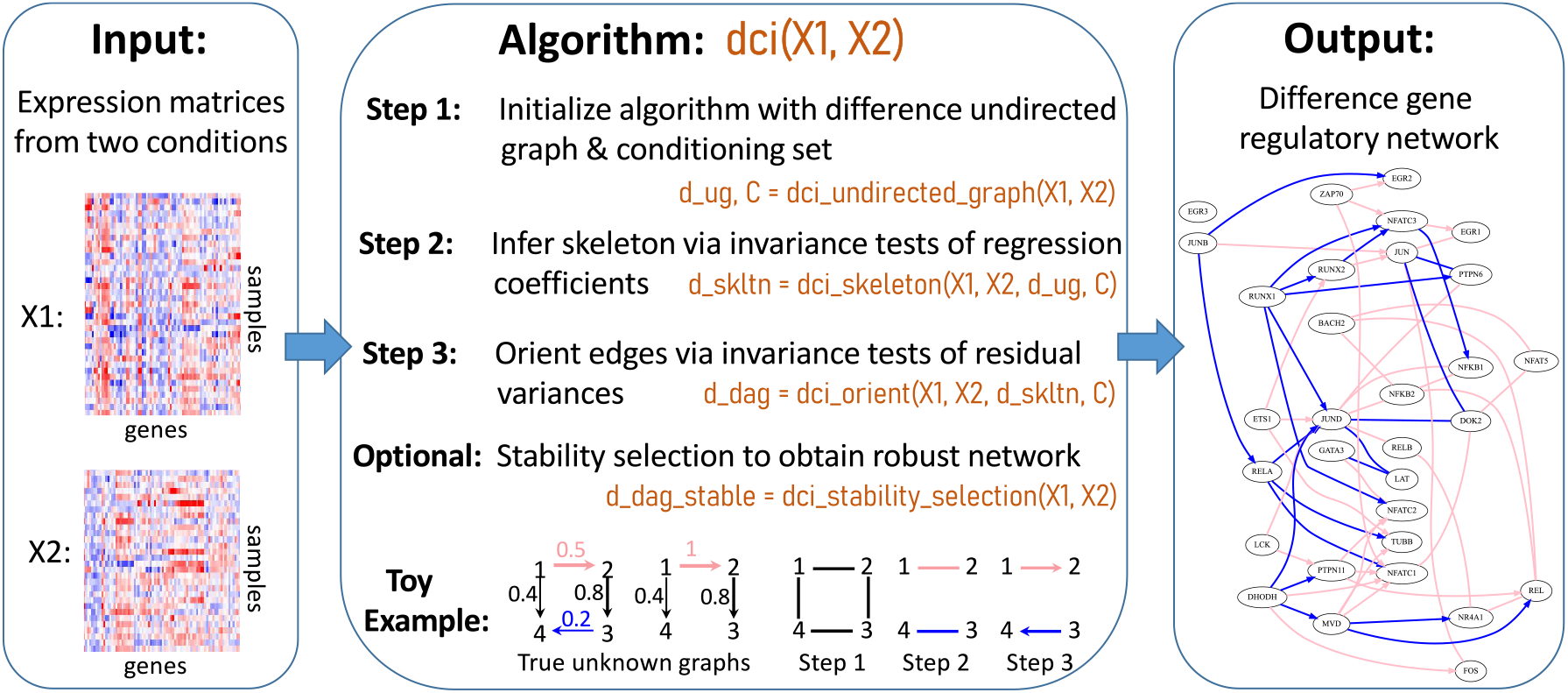
Overview of DCI algorithm: DCI takes as input two gene expression matrices *X*1 and *X*2, representing two different conditions of interest. The function `dci(X1,X2)` outputs the difference gene regulatory network consisting of the causal relationships that appeared, disappeared or changed weight between the two conditions.

The data for each condition is assumed to be generated by a linear structural equation model with Gaussian noise. More precisely, let 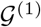 and 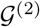 denote two DAGs on *p* nodes with weighted adjacency matrices *B*^(1)^ and *B*^(2)^. Each node *j* ∈ {1,…,*p*} in the two graphs 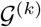, *k* ∈ {1, 2}, is associated with a random variable 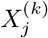, which is given by a weighted sum of its parents and independent Gaussian noise *ϵ*^(k)^, i.e.,

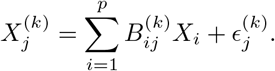

Given data 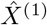 and 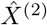 from two unknown causal graphs 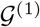 and 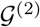, DCI determines their difference, i.e., edges *i* → *j* for which 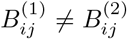. DCI consists of three steps described below (for further details see Supplementary Materials) and in Fig. 1. These steps are implemented in the `dci` function of the `causaldag` package.

### Step 1: Initialization with a difference undirected graph

To start with a reduced set of nodes and edges, DCI is initialized with a difference undirected graph, which represents changes of conditional dependencies among genes between the two conditions, as well as with a node set 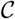, a superset of the nodes in the difference undirected graph consisting of nodes to be considered as conditioning sets in the downstream hypothesis tests. The undirected difference graph and the node set 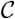 can be estimated using previous methods such as KLIEP (Liu *et al.*, 2017), which we implemented in the function `dci_undirected_graph.` Alternatively, DCI can be initialized based on a user’s prior biological knowledge or, if the number of genes to be considered is small, with the complete graph.

### Step 2: Estimation of the skeleton of the difference causal graph

The function dci_skeleton removes edges from the difference undirected graph to obtain the skeleton of the difference causal graph by testing for invariance of regression coefficients. Note that each entry 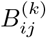 corresponds to a particular regression coefficient *β_ij|S_*, namely obtained when regressing 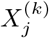 on 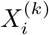 given the parents of node *j* in 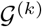. Thus, testing whether 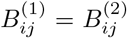 is equivalent to testing whether there exists a set of nodes 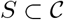 such that 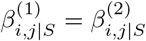. These regression coefficients are estimated from the data 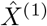 and 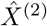 and invariance of the regression coefficients across *k* ∈ {1, 2} is tested via an F-test.

### Step 3: Orienting edges in the difference causal graph

While not all edge directions in the difference causal graph are identifiable from observational data, the function `dci_orient` orients all identifiable edges by testing for invariance of residual variances giving rise to a partially oriented difference gene regulatory network. For any edge *i* — *j* in the undirected graph obtained in Step 2, if there exists a set of nodes 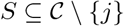 such that the residual variances satisfy 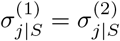, then the edge is directed as *i* → *j* if *i* ∈ *S* and *j* → *i* otherwise (see Supplementary Materials). The residual variances are again estimated from the data 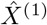 and 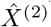 and their invariance across *k* ∈ {1, 2} is tested via an F-test.

### Stability selection to obtain robust difference gene regulatory network

DCI requires choosing hyperparameters for each step, namely the regularization parameter for KLIEP in step (1) and the significance levels for the hypothesis tests of invariance of regression coefficients in step (2) and residual variances in step (3). To overcome the difficulty of selecting hyperparameters for model selection, Meinshausen and Bühlmann (2010) proposed stability selection, which achieves family-wise error rate control and has been successfully applied for learning causal graphs in genomics (Meinshausen *et al.*, 2016). The function `dci_stability_selection` implements DCI with stability selection by running the DCI algorithm across a grid of tuning parameter combinations and bootstrap samples of the datasets 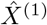 and 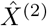. The results are aggregated and only edges with a stability score above a predefined threshold are output in the difference causal graph.

## 3 Applications and Conclusions

We applied DCI to two single-cell gene expression datasets, namely CROP-seq (Datlinger *et al.*, 2017) and Perturb-seq (Dixit *et al.*, 2016). Both datasets also contain interventional gene expression data from knockouts, thereby allowing us to assess the performance of DCI. In particular, we applied DCI to the observational single-cell data and evaluated it using an ROC curve based on the interventional data (see Supplementary Materials). By applying DCI to the CROP-seq and Perturb-seq data respectively, we learned the difference gene regulatory network between naive and activated T-cells (SI Fig. S1–S3) as well as between pre- and post-stimulation of dendritic cells with LPS (SI Fig. S4–S6). In both cases DCI outperforms the naive approach of estimating two causal graphs separately and taking their difference, and can provide valuable mechanistic insights into the biological processes of interest.

We developed the DCI package for learning differences between gene regulatory networks based on gene expression data from two different conditions of interest, such as healthy and diseased, different cell types or developmental time points. Our package is implemented in Python for ease-of-use and also includes functionality to ensure that the output difference gene regulatory network is stable and robust across different hyperparameters and data subsampling.

## Supplementary Note

### Difference Causal Inference (DCI) algorithm

In the following, we provide more details regarding the DCI algorithm; for a theoretical analysis of the algorithm see also [1]. Let 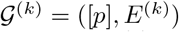 for *k* ∈ {1, 2} be a directed acyclic graph (DAG) with nodes [*p*]:= {1,…,*p*} and directed edges *E*^(*k*)^. The DAGs 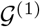 and 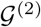 model the gene regulatory networks in the two conditions of interest. We assume that the two DAGs are consistent with the same ordering, meaning that there cannot be an edge *i* → *j* in 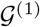 if there is a directed path *j* → ⋯ → *i* in 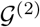 and vice-versa. This assumption is reasonable in gene regulatory networks, since genetic interactions may appear or disappear or change edge weights, but generally do not change directions. For each graph we associate a random variable 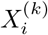 to each node *i* ∈ [*p*]. Recall that we consider the setting where we have data from two conditions and this data is generated by a linear structural equation model

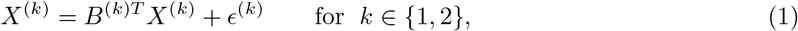

where *X* = (*X*_1_,…,*X_p_*)^*T*^ is a random vector, *B*^(*k*)^ denotes the weighted adjacency matrix of the DAG 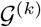 and 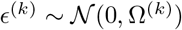 denotes Gaussian noise with covariance matrix 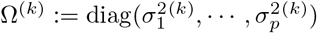. Given samples 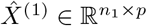 and 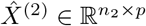 from the two models (where *n*_1_ and *n*_2_ denote the sample size under each condition), our goal is to estimate the difference-DAG across the two conditions. The difference-DAG is denoted by Δ = ([*p*], *E*) and contains an edge *i* → *j* ∈ *E* if and only if 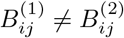.

Algorithm 1 describes the three steps of our DCI algorithm for computing the difference-DAG. In the first step, the algorithm is initialized with a difference undirected graph, which we denote by 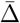, with edge *i* — *j* if and only if 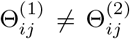 for *i* = *j*, where θ^(1)^ and θ^(2)^ are the precision matrices corresponding to the DAGs 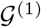 and 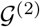. This is done to remove some edges to reduce the downstream computational burden. The difference undirected graph can be determined either using previous methods such as KLIEP [2-6], based on prior biological knowledge, or simply with the complete graph when the number of considered genes is small. In addition, to reduce the number of downstream hypothesis tests, the nodes to be considered as conditioning sets can be reduced to the nodes in the difference undirected graph as well as nodes whose conditional distribution changes between the two conditions, namely 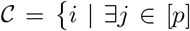 such that 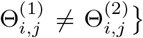. The reduced node set can be determined from the output of methods such as KLIEP [2-6], prior biological knowledge, or taken as the set of all nodes when the number of genes to be considered is small.

In the second step, the skeleton of the difference-DAG, denoted by 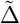, is estimated via Algorithm 2. This is done by calculating regression coefficients 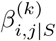 and testing whether they are invariant, i.e. whether 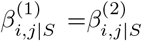, using an F-test. Given *i,j* ∈ [*p*] and *S* ⊆ [*p*] \ {*i,j*}, the regression coefficient 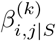 is defined as the entry in 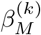 corresponding to *i*, where 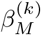 is the best linear predictor of 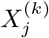 given 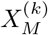, i.e., the minimizer of 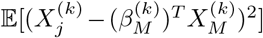 and *M*:= {*i*}∪*S*. Hence, 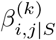 can be computed in closed form. Note that 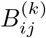 corresponds to a particular regression coefficient, namely when *S* = Pa^(*k*)^(*j*) \ {*i*}, where Pa^(*k*)^(*j*) denotes the parents of node *j* in 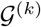. This means that we can determine whether 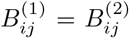 without learning each graph 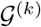, namely by testing subsets *S*: if there exists a subset *S* such that 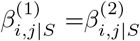, then 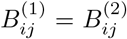 and hence the edge 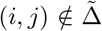. In fact, it turns out that it is sufficient to consider conditioning sets 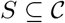 [1].

Finally, in the third step we direct edges in the skeleton of the difference-DAG 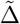 using Algorithm 3. Similar to many prominent causal inference algorithms such as the PC [7] and GES [8] algorithms, we may not be able to determine the directions of all edges in 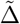, since in general, the difference-DAG Δ is not completely identifiable. In fact, we are able to identify the direction of all edges adjacent to nodes whose internal node variances are unchanged across the two conditions, i.e. for which 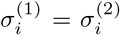 [1]. Hence the output of the DCI algorithm is a partially directed acyclic graph, which contains both directed and undirected edges. Edge directions in the difference-DAG are determined by calculating residual variances 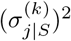 and testing whether they are invariant, i.e. whether 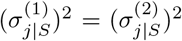, again using an F-test. Given *j* ∈ [*p*] and *S* ⊆ [*p*] \ {*j*}, the residual variance 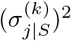 is defined as the variance of the regression residual when regressing 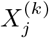 onto the random vector 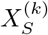. In fact it holds that 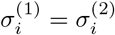 if and only if there exists a subset 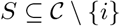 such that 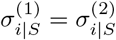 and if *i* → *j* in Δ then *j* ∈ *S*, whereas if *j* → *i* in Δ then *j* ∈ *S* [1]. Hence determining conditioning sets that lead to the invariance of residual variances can be used to orient some of the edges in the difference-DAG Δ.

**Algorithm 1.**
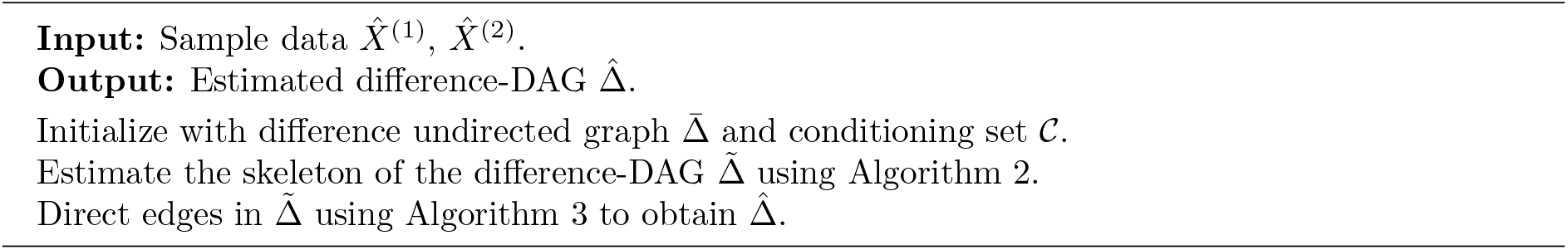
Difference Causal Inference (DCI) algorithm (`dci` function)

**Algorithm 2.**
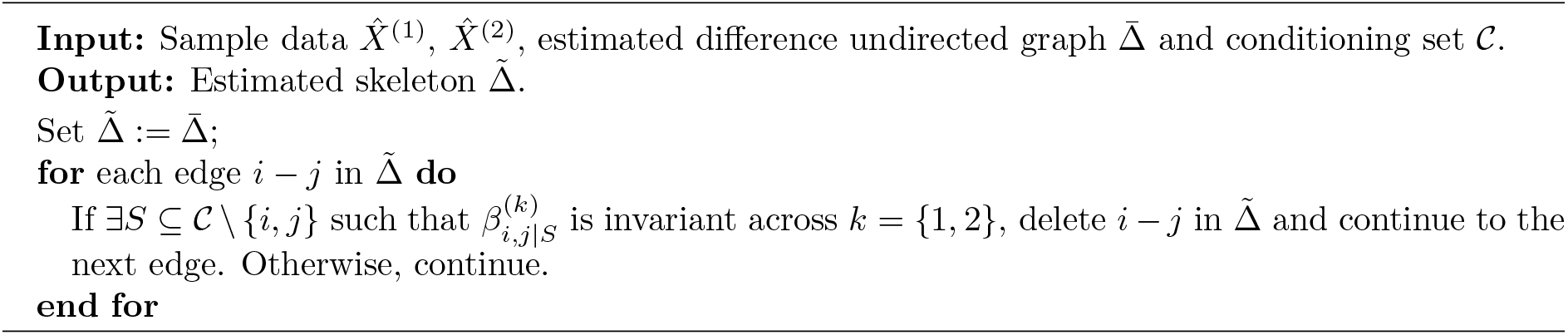
Estimating skeleton of the difference-DAG (`dci_skeleton` function)

**Algorithm 3.**
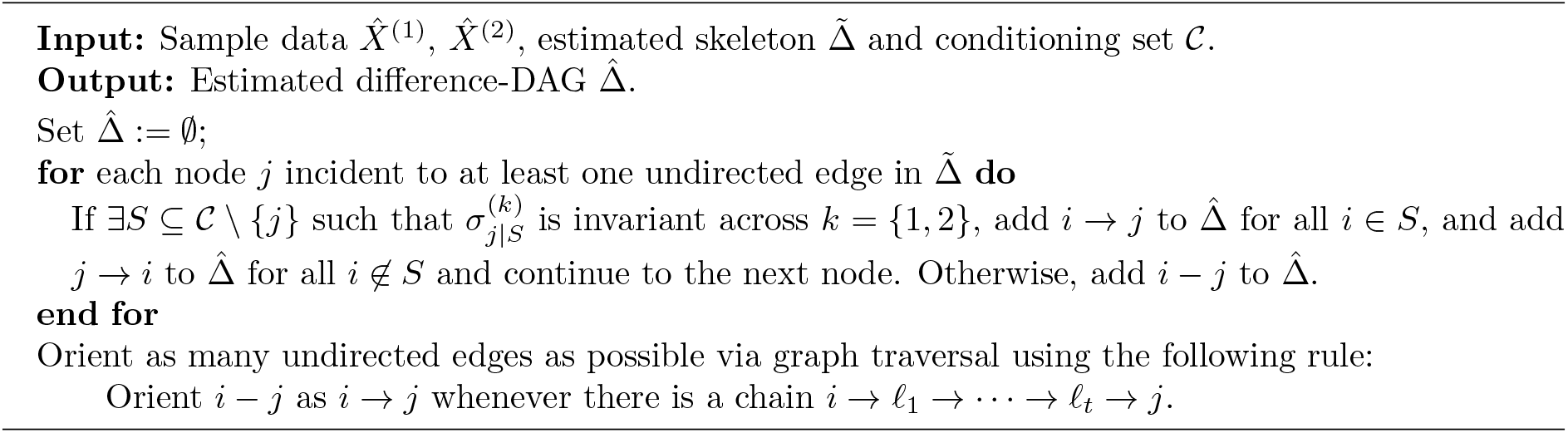
Directing edges in the difference-DAG (`dci_orient` function)

### DCI with stability selection

Running DCI requires choosing several hyperparameters, namely the *l*_1_-regularizer for estimating the difference undirected graph via KLIEP [3] as well as the significance levels for hypothesis testing of invariance of regression coefficients as well as residual variances. We implemented DCI with stability selection to address the issue of choosing the correct hyperparameters. Stability selection was introduced by [9] and has been successfully applied in tandem with other causal inference methods [10]. The idea behind stability selection is to choose the most stable estimate across different hyperparameters as opposed to focusing on choosing the right value for the hyperparameters.

Algorithm 4 outlines the methodology for running DCI with stability selection. Let Λ denote the set of considered hyperparameter values consisting of *l*_1_ regularizers for KLIEP, significance levels for hypothesis testing of invariance of regression coefficients and significance levels for hypothesis testing of invariance of residual variances. Given a particular λ ∈ Λ, we can run DCI (Algorithm 1) and obtain the corresponding estimated difference causal graph 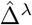. Stability selection relies on estimating the probability of selection of each edge 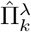 by running the DCI algorithm on subsamples of the data. Aggregating selection probabilities across different tuning parameters λ ∈ Λ, we keep edges with high selection probability as the stable set of estimated edges in the difference-DAG 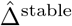.

### Evaluation on real data

We evaluate DCI for learning the causal difference gene regulatory network on single-cell gene expression data and quantify its performance in predicting the effects of gene perturbations. Note that a major advantage of our work is the ability to learn a causal as opposed to an undirected graph, which enables us to predict the effects of interventions on genes and evaluate them against true effects of interventions, measured experimentally. In the following, we assess the performance of DCI on two datasets collected via CROP-seq [11] and Perturb-seq [12]. Both of these experimental techniques collect, in a pooled fashion, single-cell gene expression data with no interventions (observational data) as well as single-cell gene expression data where some genes were knocked out via CRISPR/Cas9 (interventional data). We use the observational data to learn a causal difference gene regulatory network via DCI and evaluate this graph against the held-out CRISPR/Cas9 gene knockouts, similar in spirit to prior evaluations of causal inference methods [13].

**Algorithm 4.**
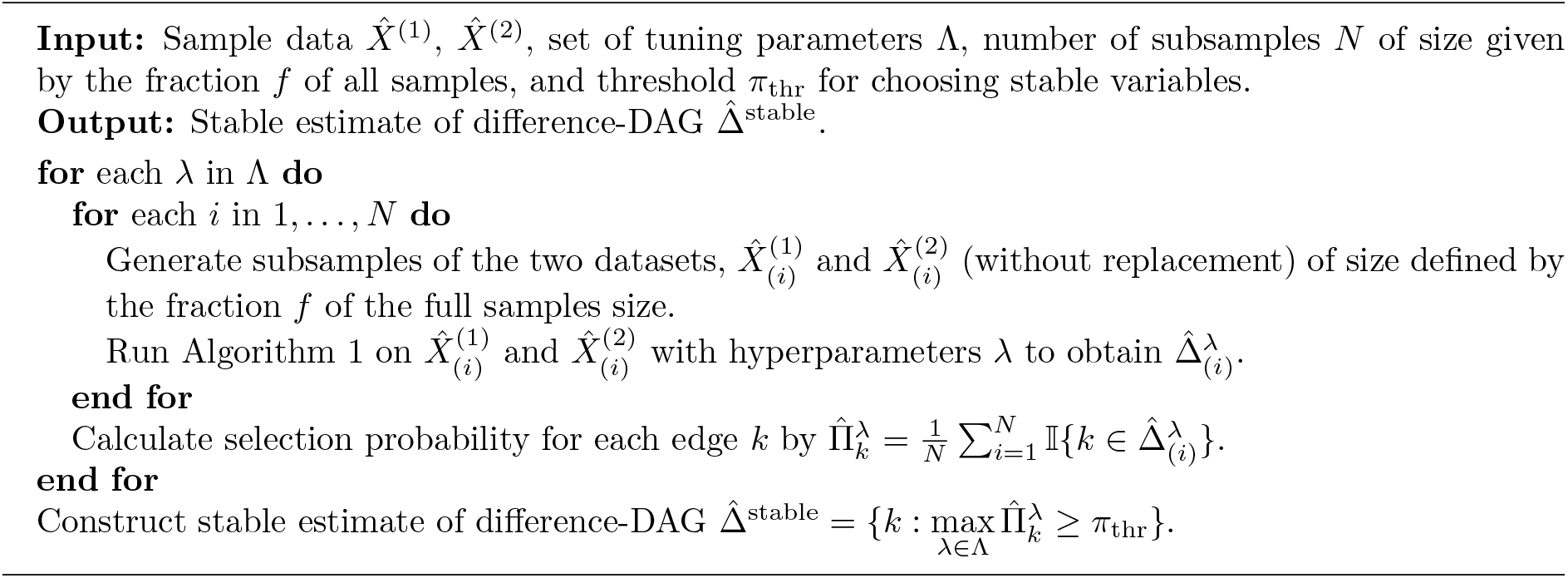
DCI with stability selection (`dci_stability_selection` function)

### CROP-seq: Naive versus activated T cells

We test our method on gene expression data collected via CROP-seq for naive and activated Jurkat T cells. In particular, we use DCI to learn the differences in the gene regulatory networks as a result of T-cell activation. The CROP-seq data includes 615 observational naive Jurkat T cells and 1320 observational activated Jurkat T cells. As in the original CROP-seq study [11], we normalize the gene expression of each cell by the total number of reads corresponding to the cell, scale expression by 10^4^ and apply a log_2_(*x* + 1) transformation to the data. The data is mean-centered prior to applying our algorithm. We follow [11] in focusing on genes most relevant to T-cell activation and keep genes that have non-zero variance, resulting in 31 genes.

We apply DCI on the observational naive and activated gene expression data to directly obtain the causal difference gene regulatory network (difference-DAG), which contains edges that appeared, disappeared or changed weight between the two cell states. We report the performance of DCI when initialized in the complete graph as well as when initialized with the difference undirected graph estimated via KLIEP (*l*_1_ regularization set to 0.005). Additionally, we compared the performance of DCI to the naive approach of running classical causal inference algorithms such as PC [7] or GES [8] on each dataset (naive and activated) separately, obtaining two causal graphs and then taking the difference. We consider an edge to be in the difference-DAG if the edge was directed in one causal graph and absent in the other causal graph.

As previously mentioned, we can use gene knockouts, collected as part of the CROP-seq study for evaluation of the causal difference gene regulatory network. Note that if perturbing a gene affected the gene expression distribution of another gene, this means that the perturbed gene is upstream of the affected gene in the gene regulatory network. In the following we describe how we estimate the differences in the effects of CRISPR/Cas9 perturbations on genes between the two states (naive and activated) to construct an ROC curve for evaluating the DCI algorithm versus naive applications of PC and GES.

First, for each condition (naive and activated), we separately obtain a matrix that describes which gene knockouts had an effect on which genes (Figures S1a and S1b). Then, we take the difference between these matrices to determine the differences in the effects of perturbations (Figure S1c). In order to construct the matrices in Figures S1a and S1b, for each condition, we estimate the impact of each gene deletion *j* ∈ {1,…, *d*} on each of the measured genes *i* ∈ {1,…,*p*} by testing whether the observational distribution (no intervention) of the measured gene *i* is significantly different from the interventional distribution of the measured gene *i* when gene *j* was deleted using a Wilcoxon rank-sum test. We form a *p* × *d* matrix of p-values, *Q*, from the Wilcoxon rank-sum tests. Next, each column *j* in *Q* is thresholded using the entry *q_jj_*, which is the p-value obtained by comparing the distribution of the gene expression level of a deleted gene versus its distribution without intervention. The rationale is that knocking out a particular gene should result in a change in its own gene expression distribution and can be used as a baseline to threshold the other entries in the column. In particular, we conclude that *q_ij_* is significant if and only if *q_ij_*≤*q_jj_*. After thresholding the matrix *Q* in this manner, we obtain the binary matrices in Figures S1a and S1b, which summarize the effects of the interventions. By forming the difference of these binary matrices we obtain the binary matrix *Q*^Δ^ in Figure S1c. Since not all CRISPR/Cas9 knockouts were effective, here we focused our analysis on the top most effective interventions, which were prioritized based on the maximum *q_jj_* p-value (taken over two conditions), using the mean p-value as the cutoff to filter interventions.

We use the matrix of differences in the effects of interventions to evaluate DCI, PC and GES by constructing an ROC curve. If the predicted difference-DAG has a directed edge from *j* to *i*, we count this edge as a true positive if 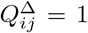, i.e. there was a difference in the effect of knocking out gene *j* on gene *i* between the two conditions. If the predicted difference-DAG has a directed edge from *j* → *i* but 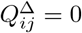, the edge is counted as a false positive. Note that this definition of a false positive is overly conservative, since we may have 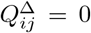 if *q_jj_* is significant in both matrices, but the magnitude of the effect changes. In other words, 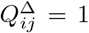 only captures additions/deletions of edges, but does not capture changes in edge weights. We construct an ROC curve by varying the parameters of DCI, PC and GES. The ROC curve in Figure S2 shows that DCI outperforms PC and GES in predicting the effects of interventions on this single-cell gene expression dataset. In Figure S3, we include examples of the estimated difference gene regulatory networks inferred via DCI (our algorithm) and GES (the best performing baseline).

### Perturb-seq: Dendritic cells at 0 versus 3 hours post-stimulation

We perform a similar evaluation of DCI on gene expression data collected as part of the Perturb-seq dataset [12]. Gene expression data was collected from bone-marrow derived dendritic cells (BMDCs) prestimulation (0 hours) and after stimulation with LPS (3 hours). We applied DCI to learn the difference gene regulatory network between these two time points. We used the same procedure for pre-processing Perturb-seq data as we used for CROP-seq. Additionally, we filtered cells for quality, only keeping cells with at least two nonzero counts (CROP-seq dataset already satisfied this filtering constraint). The filtered Perturb-seq data includes 940 observational cells collected at 0 hours and 990 observational cells collected at 3 hours. We followed [12] in focusing on 24 transcription factors that are important for dendritic cell regulation.

Using the same procedure as performed on the CROP-seq dataset, we constructed the binary matrices describing the effects of gene deletions on measured genes for the two time points (0 and 3 hours) separately, shown in Figures S4a and S4b, and then determined the difference in the effects of the interventions between the two time points in Figure S4c. As above, we constructed an ROC curve, taking the differences in the effects of interventions as the ground truth. The ROC curve (Figure S5) shows that in the majority of settings, DCI outperforms the naive approach of estimating two causal graphs separately via PC or GES and taking the difference of the output graph. Figure S6 shows examples of the estimated difference gene regulatory networks inferred via DCI (our algorithm) and GES (best performing baseline).

---

**Fig. S1:**
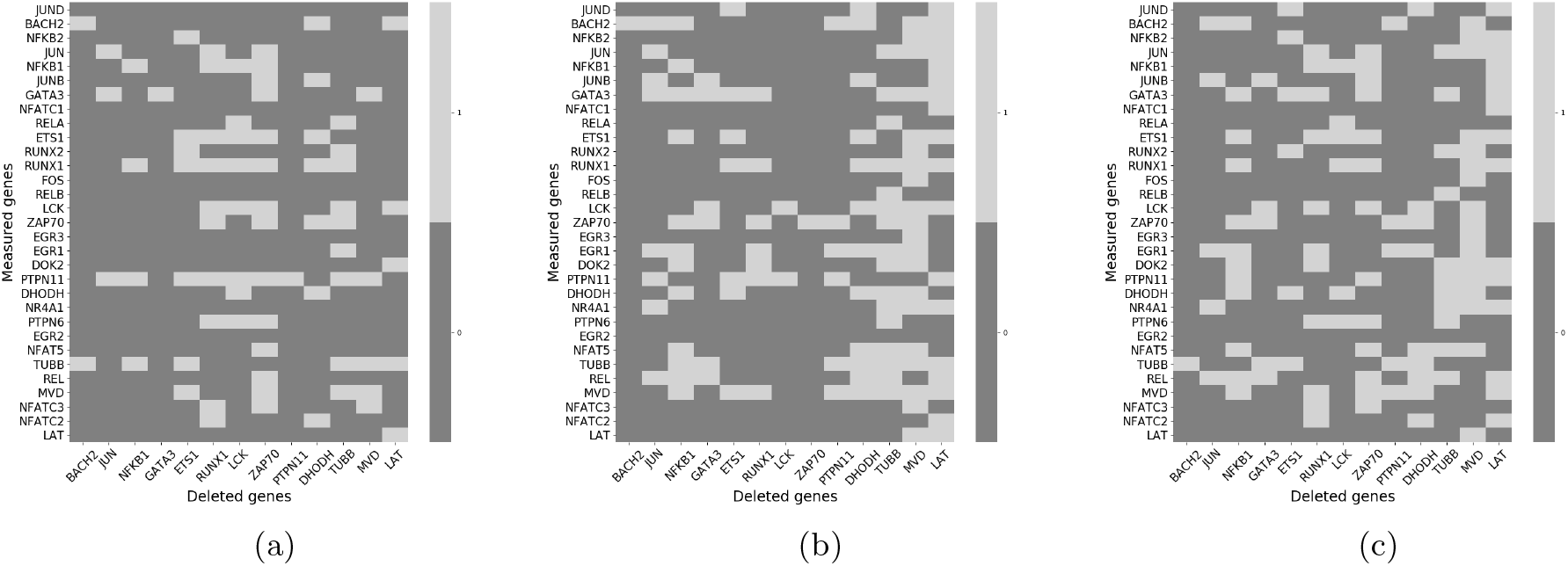
Effects of gene deletions estimated from CROP-seq data; (a) naive T cells, (b) activated T cells, and (c) the difference between the binary matrices in (a) and (b), i.e., the difference in the effects of each gene deletion on the measured genes between naive and activated T cells; this binary matrix is taken to be the ground truth for constructing ROC curves.

**Fig. S2:**
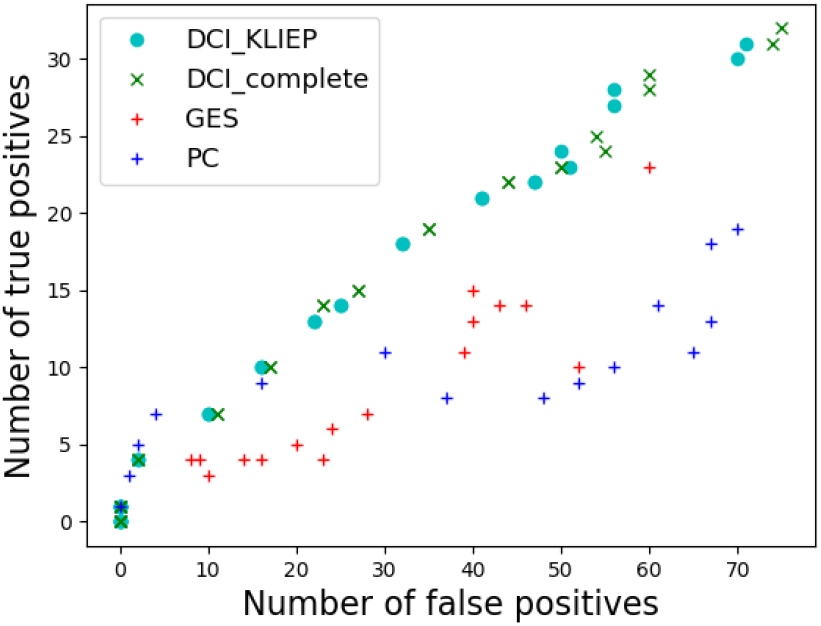
ROC plot evaluating DCI (initialized in the undirected difference graph estimated via KLIEP as well as in the complete graph), GES and PC on the CROP-seq data for predicting the differences in the effects of gene knockouts.

**Fig. S3:**
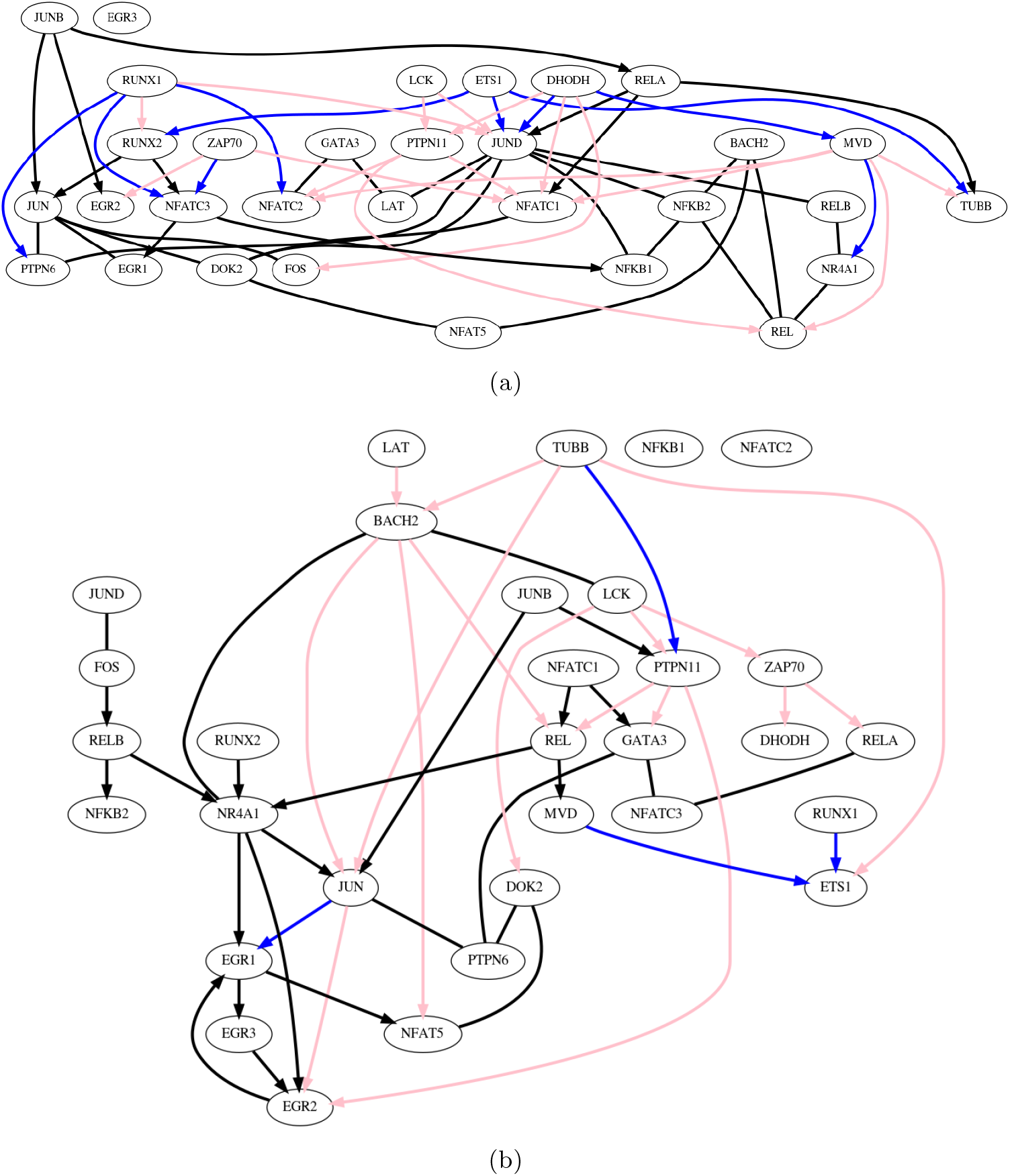
Examples of difference gene regulatory networks between naive and activated Jurkat T cells, estimated from the CROP-seq data. Difference gene regulatory network inferred via (a) our algorithm, DCI, initialized with KLIEP, which directly learns the difference causal graph from two datasets and (b) baseline causal structure discovery algorithm, GES, which estimates two gene regulatory networks separately and then takes the difference. Blue edges indicate true positives and pink edges indicate false positives. Black edges are the edges inferred to be in the difference gene regulatory network for which ground truth is not available. Graphs were chosen such that the number of false positives is the same across the two methods (16 false positives).

**Fig. S4:**
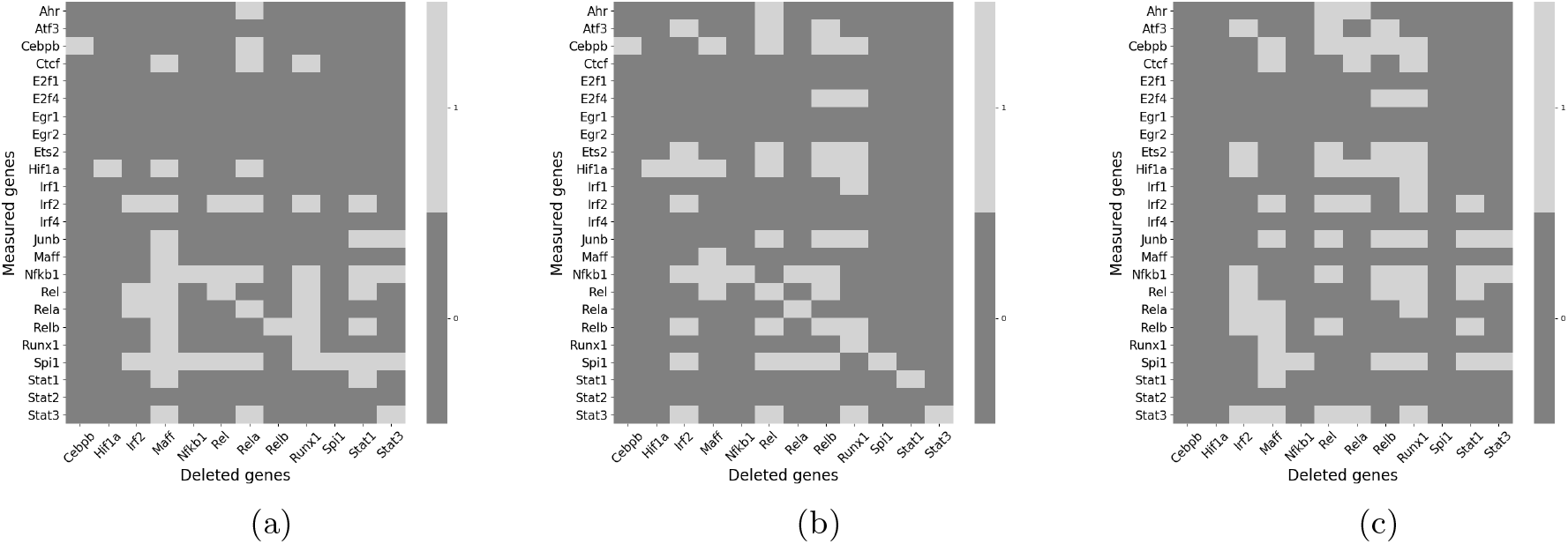
Effects of gene deletions estimated from Perturb-seq data; (a) before stimulation with LPS, (b) after stimulation with LPS, and (c) the difference between the binary matrices in (a) and (b), i.e., the difference in the effects of each gene deletion on the measured genes before and after stimulation with LPS; this binary matrix is taken to be the ground truth for constructing ROC curves.

**Fig. S5:**
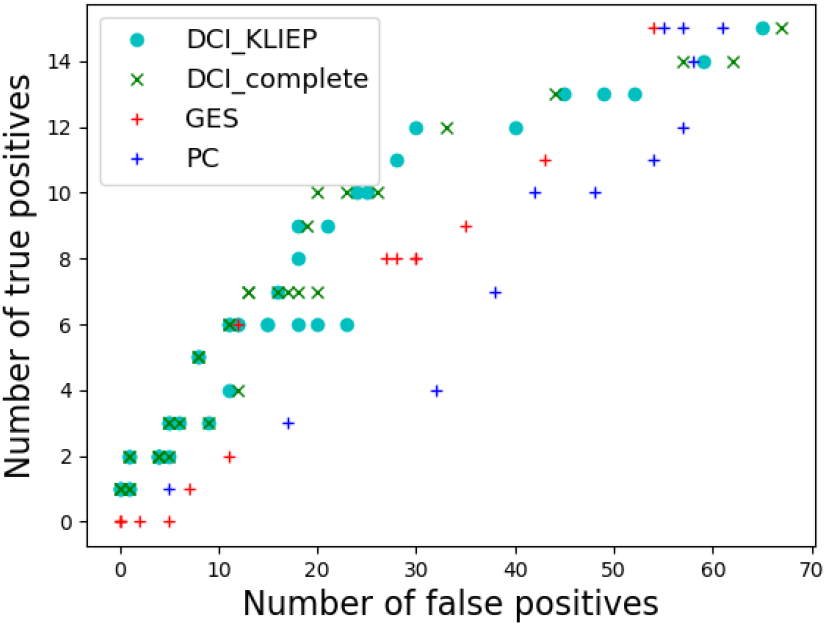
ROC plot evaluating DCI (initialized in the undirected difference graph estimated via KLIEP as well as in the complete graph), GES and PC on the Perturb-seq data for predicting the differences in the effects of gene knockouts.

**Fig. S6:**
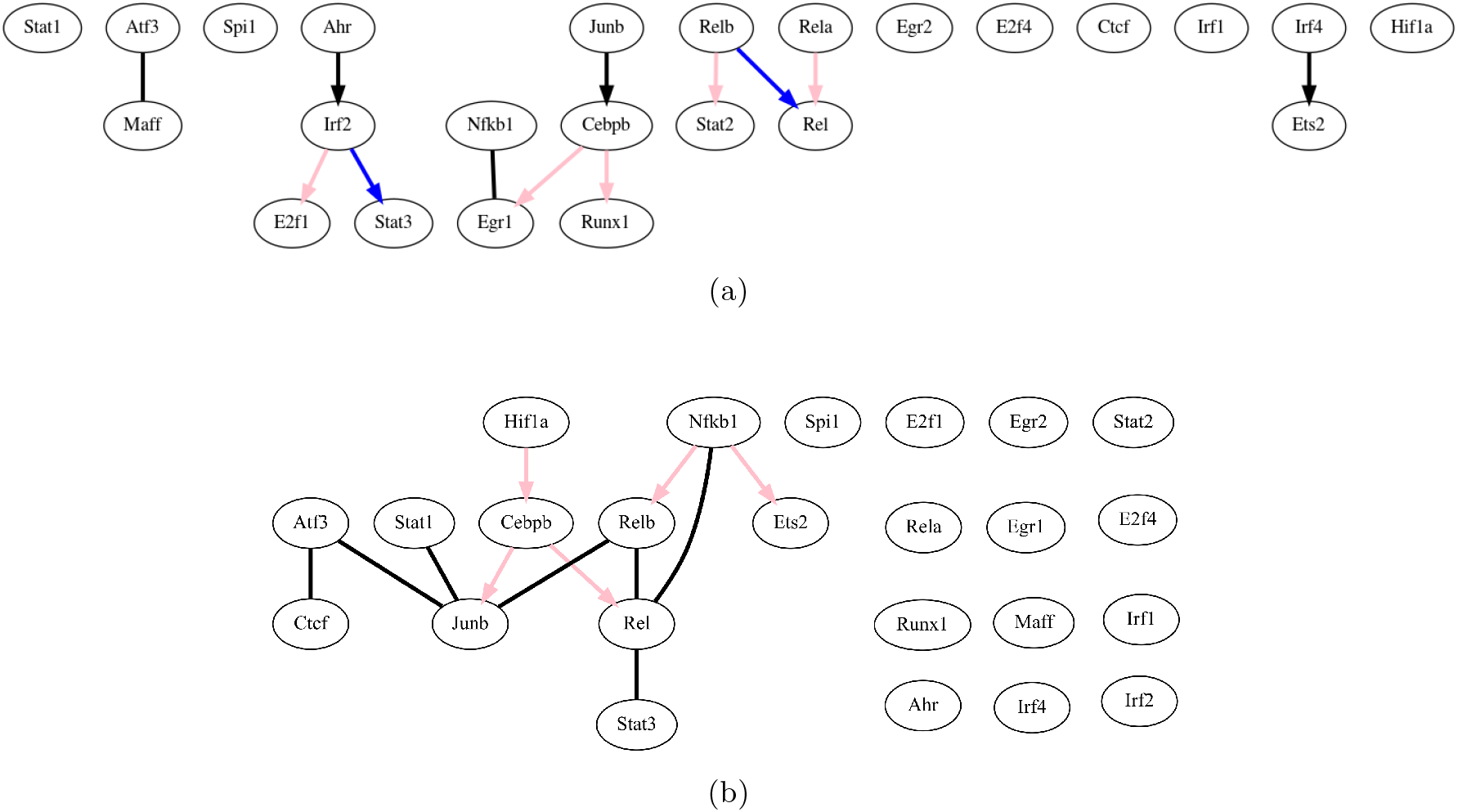
Examples of difference gene regulatory networks of dendritic cells before and after stimulation with LPS, estimated from the Perturb-seq data. Difference gene regulatory network inferred via (a) our algorithm, DCI, initialized with KLIEP, which directly learns the difference causal graph from two datasets and (b) baseline causal structure discovery algorithm, GES, which estimates two gene regulatory networks separately and then takes the difference. Blue edges indicate true positives and pink edges indicate false positives. Black edges are the edges inferred to be in the difference gene regulatory network for which ground truth is not available. Graphs were chosen such that the number of false positives is the same across the two methods (5 false positives).

